# *In vivo* generation of DNA sequence diversity for cellular barcoding

**DOI:** 10.1101/004499

**Authors:** Ian D. Peikon, Diana I. Gizatullina, Anthony M. Zador

## Abstract

Heterogeneity is a ubiquitous feature of biological systems. A complete understanding of such systems requires a method for uniquely identifying and tracking individual components and their interactions with each other. We have developed a novel method of uniquely tagging individual cells *in vivo* with a genetic “barcode” that can be recovered by DNA sequencing. We demonstrate the feasibility of this technique in bacterial cells. This method should prove useful in tracking interactions of cells within a network, and/or heterogeneity within complex biological samples.

## INTRODUCTION

Reverse engineering any complex system requires the simultaneous monitoring of individual components. Recent advances in high-throughput DNA sequencing have given biologists unprecedented access to massively parallel data streams. Genetic barcoding - the labelling of individual cells with a unique DNA sequence - when combined with these technologies, will enable monitoring of millions or billions of cells within complex populations. This approach has proved useful in tagging neurons (1)4/23/14 11:02 PM and hematopoietic stem cells (2,3) for lineage analysis and could be applied to the normal and/or abnormal development of other cell populations or tissues, including tumours. The simultaneous monitoring of millions or billions of bacterial or yeast cells within a population will allow a detailed analysis of population genetics and growth dynamics under various conditions. Moreover, interactions between cells within cellular networks can be uncovered by passing barcodes between interacting cells (i.e. B- and T-cells, neurons, tumours, etc.). Indeed, *in vivo* barcoding of individual neurons is the requisite first step towards converting neuronal connectivity into a form readable by high throughput DNA sequencing (4), which is currently a major effort in our lab.

Most current approaches for tagging individual cells with a genetic barcode rely on the creation of diverse libraries in vitro and subsequent delivery of genetic material into a host cell at low-copy number. Such in vitro approaches are limited by cloning bottlenecks that cause reduced library diversities and sequence biases, by incomplete labelling of all cells within a population, by the possibility of introducing multiple barcodes per cell, and by the challenges of working across organisms (e.g. retroviral barcoding cannot be applied in some organisms like *C. elegans*).

*In vivo* barcoding has the potential to overcome all of these limitations. Mechanisms for generating diversity *in vivo* exist, endogenously, in many organisms - most notably the mammalian immune system. However, efforts to repurpose the immune system’s V(D)J recombination for *in vivo* cellular barcoding (5) yielded limited barcode diversity - on the order of a dozen unique sequences - in cells other than lymphocytes (6). Exogenous recombinases have been successfully applied to generate diverse combinations of colors for cellular tagging purposes. This technique, better known as Brainbow (7), relies on Cre recombinase to rearrange a cassette resulting in the stochastic expression of a subset of different colored fluorescent proteins (XFPs) in neurons. The theoretical diversity of Brainbow is in the hundreds, but cannot be easily assayed with DNA sequencing because it relies heavily on gene copy number variation as well as recombination. We reasoned that by replacing XFPs with unique sequences, we could design a barcoding system with the potential to achieve diversities that matched the scale of high-throughput sequencing technologies.

We have developed a novel method of generating sequence diversity *in vivo* for the purpose of cellular barcoding. Our method, which relies on a DNA invertase - Rci, recombinase for clustered inversion (8) - to shuffle fragments of DNA, has the potential to easily achieve diversities over 10^9^ unique sequences. Here, we show that this method can be applied for the *in vivo* generation of diversity in bacterial cells.

## MATERIAL AND METHODS

### In silico simulations

We performed *in silico* simulations to determine the behaviour of different cassette architectures. For Cre-based cassettes, n_cell_ cassettes (n_cell_=10000) of n_frag_ fragments (n_frag_=100) were operated on independently. Each fragment was flanked on its 5’ end with a *loxP* site in sense orientation (5’-GCATACAT-3’) and on it’s 3’ end with a *loxP* site in the antisense orientation. Concatenation of fragments resulted in cassettes in which adjacent fragments (excluding ends) were separated by two *loxP* sites in opposing orientation. We defined Cre recombination as two independent binding events to *loxP* sites. Binding of Cre to a pair of *loxP* sites always resulted in recombination, where the result (inversion or excision) was defined by the relative orientation of the sites defined in a look up table (updated after each event). Completion is defined to be the point at which Cre can no longer mediate an excision event. The number of recombination events required to reach completion was tracked for each cassette. For Rci-based cassettes, n_cell_ cassettes (n_cell_=10000) of n_frag_ fragments (n_frag_=10) were operated on independently. Here, we considered two architectures. For the first architecture, each fragment of the cassette was flanked on its 5’ end with an *sfx* site in sense orientation and on its 3’ end with an *sfx* site in the antisense orientation. Concatenation of fragments resulted in cassettes in which adjacent fragments (excluding ends) were separated by two *sfx* sites in opposing orientation. For the purpose of simulations, we consider each pair of *sfx* sites between fragments to be equivalent to a single bidirectional *sfx* site. We defined Rci recombination as two independent binding events to *sfx* sites. In this case, binding of Rci to a pair of *sfx* sites always resulted in recombination (inversion). Simulations were allowed to proceed for *m* recombination events per cassette. For the second Rci architecture, the 5’ end of the cassette begins with a single *sfx* site in sense orientation, followed by a single sequence fragment. The cassette is extended by addition of an *sfx* site and a sequence fragment, with the orientation of *sfx* sites alternating throughout the cassette. The cassette is terminated at its 3’ end by an *sfx* site in antisense orientation. We defined Rci recombination as two independent binding events to *sfx* sites. Binding of Rci to a pair of *sfx* sites only resulted in recombination if the *sfx* sites were in opposite orientations (inversion only). Simulations were allowed to proceed for *m* recombination events per cassette. The code for running all simulations is provided in Supplementary materials.

### Synthesis of barcode cassettes

A 5-fragment barcode cassette was synthesized as a minigene by IDT and inserted into a standard plasmid (IDP190). Several different *sfx* sites were used to avoid perfect inverted repeats to simplify DNA synthesis. The cassette was constructed as ANCHOR105-*sfx101R*-BC1-*sfx102L*-BC2-*sfx106R*-BC3-*sfx112L*-BC4-*sfx109R*-BC5-*sfx101L*-ANCHOR56 where R indicates the *sfx* sequence in 5’-3’ orientation and L indicates the reverse orientation (where orientation is determined by the core sequence: 5’-GTGCCAA-3’). The barcode cassette was synthesized with flanking sequences for use in PCR amplification and sequencing (#3:ANCHOR105 and #4:ANCHOR56). In addition, the cassette was flanked by restriction sites (PciI) and flanking primer sequences (#1:BC_F and #2:BC_R) for subsequent PCR and cloning. An additional BC cassette (containing BC6-11) was synthesized by GeneWiz (BCextension). The cassette was constructed as BamHI-BC6-*sfx101R*-BC7-*sfx102L*-BC8-*sfx106R*-BC9-*sfx112L*-BC10-*sfx109R*-BC5-*sfx101L*-NheI.

### Plasmid construction

The 5-fragment barcode cassette (from plasmid #IDP190) was amplified using primers #1:BC_F and #2:BC_R, digested with PciI, and cloned into pet22b. A strain of E. coli harbouring the pEK204 plasmid, which encodes Rci recombinase (NCBI Reference Sequence: NC_013120.1), was ordered from NCTC (NCTC 13452: J53-derived E. coli. GenBank accession number EU935740). The open reading frame of Rci was obtained by PCR of the pEK204 plasmid with primers #7:NdeI-Rci_F and #8:NotI-Rci_R, which add restriction sites NdeI and NotI, respectively. Rci was cloned into plasmid pet22b using restriction sites NdeI and NotI (thus removing the periplasmic localization signal of pet22b) to create Plasmid #IDP205:(T7->Rci; 5BC). The constitutively active promoter, pKat (Registry of Standard Biological Parts: BBa_I14034, http://parts.igem.org/Part:BBa_I14034), and a ribosomal binding site (RBS): AGGAGG, flanked by restriction sites BglII and NdeI were synthesized as complementary oligos #9:s-pKat_promoter and #10:as-pKat_promoter, annealed, digested, and subsequently cloned into #IDP205 to make plasmid #DIG35:(pKat->Rci, 5BC). To extend the BC cassette we amplified #IDP205 using primers #11:BamHI-BC5_F and #12: NheI-BC5_R, and cut with BamHI and NheI. The insert, BC6-BC11, was digested from BCextension with BamHI and NheI. The backbone and insert were ligated to make plasmid #DIG70:(T7->Rci, 5BC). The constitutively active promoter, pKat, and a ribosomal binding site (RBS) flanked by restriction sites BglII and NdeI were synthesized as complementary oligos #9:s-pKat_promoter and #10:as-pKat_promoter, annealed, digested, and subsequently cloned into #DIG70 to make plasmid #DIG71:(pKat->Rci, 11BC). All cloning was performed using Top10 chemically competent cells (Invitrogen) with growth at 37degC.

### Bacterial culture & shuffling

For initial tests with T7 induced expression of Rci, plasmids IDP205 or DIG35 were transformed into BL21(DE3) bacterial cells (NEB) and grown in 5ml of normal or OvernightExpress (Millipore) supplemented media overnight. Plasmid DNA was isolated and the Rci coding sequence was removed (to prevent further shuffling) by double digestion (NdeI-NotI), blunting (Mung Bean), and religation (Roche Rapid Ligation kit). The transformed ligation was plated for clonal analysis. Clonal analysis involved the selection of single colonies, growth in LB for 16 hours, plasmid isolation, and Sanger sequencing with ANCHOR105 used as a primer. For tests of the pKat driven expression, plasmids IDP205 or DIG35 were transformed into Top10 bacterial cells (Invitrogen) and grown in 5ml of LB overnight. Plasmid DNA was isolated and the Rci coding sequence was removed (to prevent further shuffling) by double digestion (NdeI-NotI), blunting (Mung Bean), and re-ligation (Roche Rapid Ligation kit). The transformed ligation was plated for clonal analysis. Clonal analysis involved the selection of single colonies, growth in LB for 16 hours, plasmid isolation, and Sanger sequencing with ANCHOR105 used as a primer. For high-throughput sequencing by PacBio, plasmids IDP205, DIG35, and DIG71 were transformed into Top10 cells (Invitrogen) and grown overnight in 50mL of LB. Plasmid DNA was isolated and digested with PciI to release the barcode cassette. The barcode cassette was prepared for PacBio sequencing using the PacBio SMRTbell Template Prep Kit according to the manufacturer’s instructions and cassettes from each original plasmid were sequenced on a single SMRT cell. PacBio sequences were collapsed into circular consensus reads using the PacBio command line tools.

### Barcode Reconstruction

Our sequence alignment algorithm is written in Matlab and uses the Matlab Bioinformatics Toolbox. Sequencing reads are processed independently. Each known barcode fragment is aligned to the sequence read (in both orientations) with a thresholded Smith-Waterman alignment (9) and the position of the segment along the read is stored. The threshold is set by a bootstrap method. Briefly, one hundred randomly generated 100-mers are aligned to all of the sequence traces and a score is associated with each alignment. The mean score plus two standard deviations is considered the threshold for all subsequent alignments. The complete barcode can be reconstructed based on the positions of each segment. Because the algorithm relies only on local alignment, this method is extremely robust to sequencing errors.

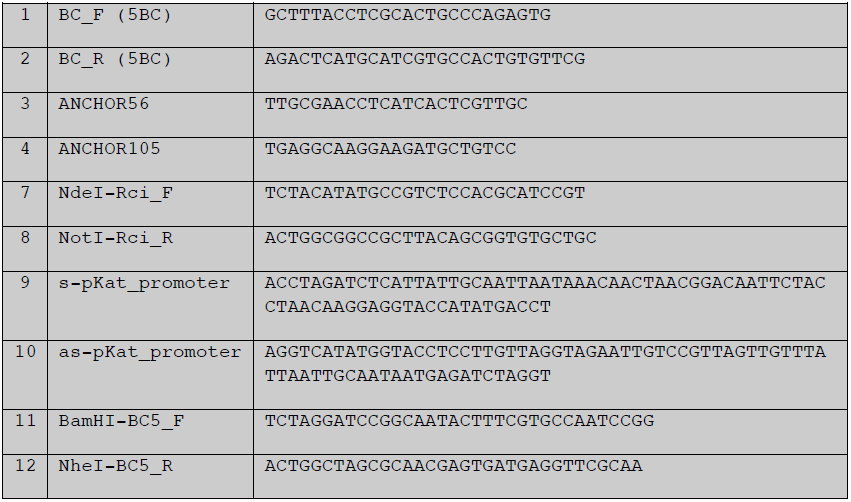
Primer Table.

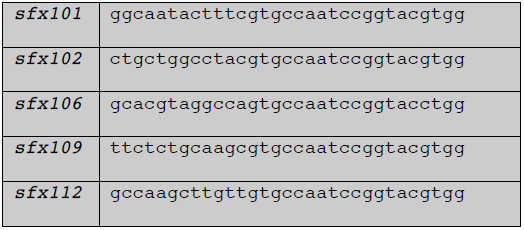
*sfx* sites used.

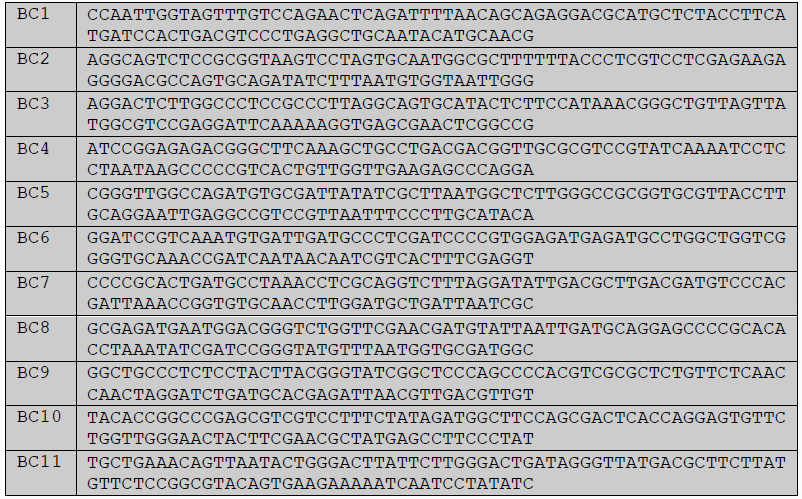
BC sequences.

Plasmid Table.

Plasmid Table
|  |  |
| --- | --- |
| IDP190 | 5BC Cassette |
| IDP205 | T7->Rci; 5BC Cassette |
| DIG35 | pKAT->Rci; 5BC Cassette |
| BCext | BC6-BC11 extension |
| DIG70 | T7->Rci; 11BC Cassette |
| DIG71 | T7->Rci; 11BC Cassette |

### Sequences of BC cassettes and Rci ORF

All relevant sequences and plasmid maps are provided in Supplementary Materials.

## RESULTS

### Design of Cre-based barcoding

Our design goals were to create a modular genetically encoded barcode system that is easily scalable, cross-platform (applicable across model organisms), compatible with high-throughput sequencing technologies, and robust to sequencing errors. Our lab is interested in generating unique barcodes to label all of the cells of the mouse cortex - approximately 1×10^7^ neurons (10). In general, if the repertoire of possible barcodes is substantially greater than the number of cells in the population of interest, then even randomly generated barcodes will label most cells uniquely. Specifically, if *n* is the number of cells and *k* is the number of possible barcodes, then under simple assumptions the fraction of uniquely labeled cells will be *e^-n/k^*. Thus assuming one barcode per cell, a barcode repertoire exceeding the number of cells by a factor of 100 will yield 99% uniquely labeled cells. Therefore, we sought a barcode system with the potential to scale to at least 100*10^7^ = 10^9^ unique barcodes.

**Figure 1.**
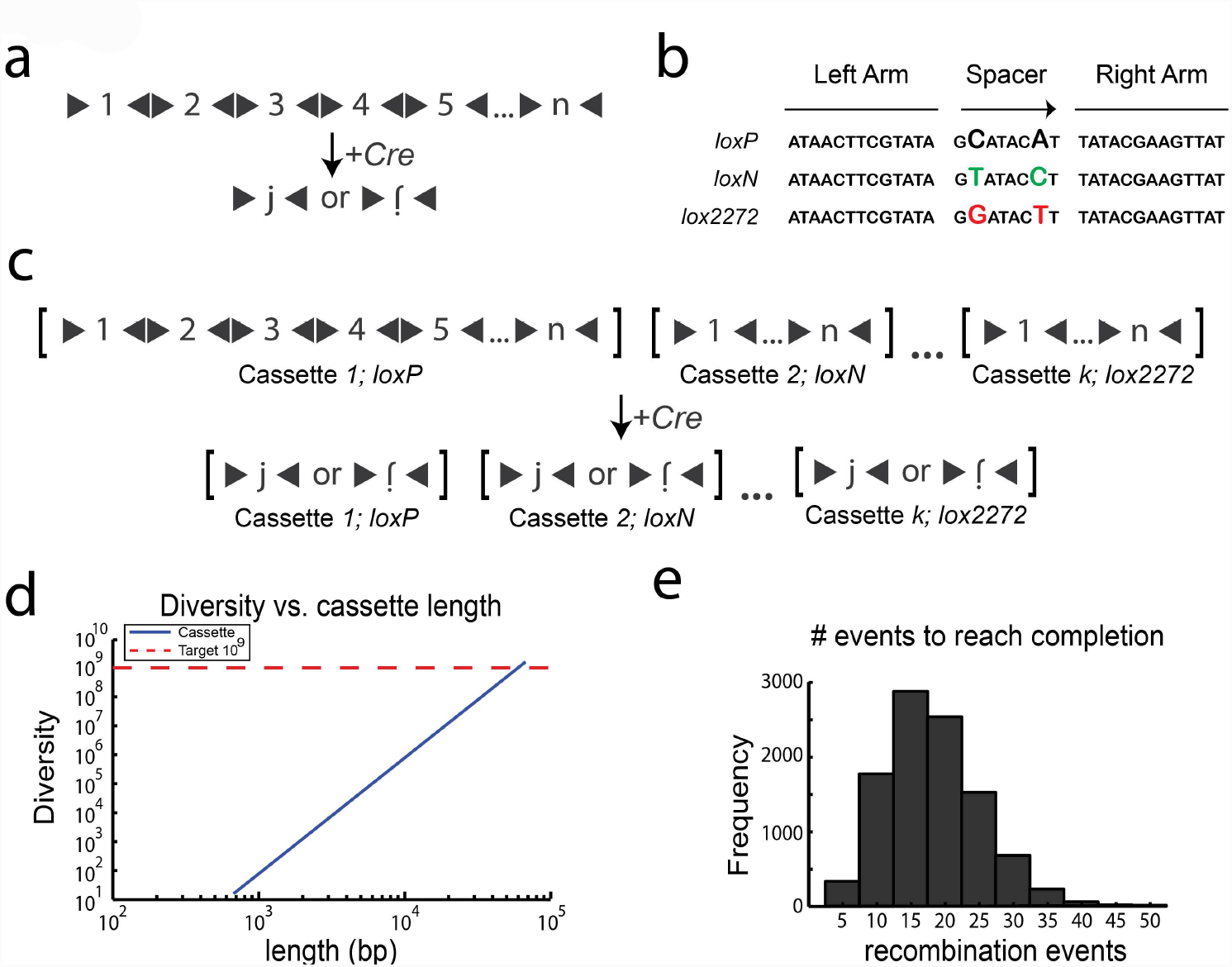
Design of Cre cassettes. **(A)** *n* sequence fragments, each flanked by *loxP* sites (black triangles) in opposing orientations are concatenated to form a cassette. Upon Cre recombination, the cassette is shuffled and pared down until a single fragment, *j*, remains in either orientation. **(B)** Several variant mutually incompatible *lox* sites have been described, each of which has a pair of mutations in the spacer sequence. **(C)** The diversity attainable by Cre recombination can be increased by concatenating *k* cassettes, where each cassette utilizes an incompatible *lox* site. Cre recombination results in the *independent* shuffling of each of the *k* cassettes. **(D)** The diversity as a function of length for cassettes containing *n=100* fragments (100bp each) concatenated together. Lengths >60kb are needed to reach the requisite diversity of 10^9^. **(E)** The distribution of the number of recombination events needed for each cassette to reach completion.

Our initial design relied on Cre recombinase to shuffle and pare down a cassette of *n* barcode fragments (each fragment flanked by *lox* sites in alternating orientation) by stochastic inversion and excision events to leave a single fragment (Figure 1a). Cre acts by binding to and mediating recombination between two of its cognate DNA sequences, called *lox* sites. Cre-mediated recombination between any two compatible *lox* sites in the same orientation causes the excision of the intervening sequence elements. In contrast, recombination between *lox* sites in opposing orientation causes the inversion of any intervening sequence elements.

This architecture (shown in Figure 1a) has a theoretical diversity of *2n* after completion, since any fragment, *j*, of the *n* fragments can end in either the forward or inverted orientation (Figure 1a). Several variant *lox* sites have been discovered with a wide variety of characteristics (11–13). The Brainbow (7) system, for example, employed three lox sites that were shown to be mutually incompatible including *loxP* (the original lox site), *loxN* (7), and *lox2272* (14) (Figure 1b). We reasoned that a Cre-based barcoding approach could be extended to achieve higher diversities by concatenating *k* cassettes of *n* fragments, where each cassette employs one of a subset of incompatible lox sites (Figure 1c). Here, the theoretical diversity is *(2n)^k^* because each cassette operates independently. Thus with *n*=100, *k*=4, theoretical diversities reach our goal of 10^9^ unique sequences.

Unfortunately, many of the reported *lox* variants have not been validated for complete incompatibility or pairwise efficiency. Moreover, because of the repetitive nature of *lox* sites it would be difficult to synthesize cassettes with the hundreds of fragments required to achieve our target diversity. For example, an architecture with *n*=100, *k*=4, where each fragment is above the minimum length (∼100bp) for efficient recombination (15), requires a large genomic insertion with a length greater than 60kb (Figure 1d). Finally, simulations suggested that Cre-based architectures are subject to considerable biases that limit the diversities that can be achieved in practice (Supplementary Note 1, Supplementary Figure 1).

### Employing DNA invertases for cassette shuffling

The key limitation of Cre-based designs is that Cre mediates both inversion and excision/insertion. Because insertion is a bimolecular reaction whereas excision is a unimolecular reaction, in general equilibrium will favour excision over insertion. Thus the equilibrium diversity of a Cre-based cassette scales linearly with the number of fragments *n*. In simulations (See Methods), 10 to 40 recombination events were performed on each cassette before reaching completion (Figure 1e). The diversity of the Cre-based cassettes could, in principle, be increased by preventing the reaction from proceeding to completion. In the limit, if only inversions were permitted, then the fragments of the cassette would be shuffled rather than pared down. Intuitively, the advantage of eliminating excision events can be understood by analogy to a deck of *n* playing cards, in which each card can occur in either orientation (face up or face down). If excisions dominate, then the diversity is given by eliminating all but one card. If inversions dominate, then the diversity is given by all the possible sequences of n shuffled cards (n!) multiplied by all of the possible orientations (2^n^). Thus eliminating excisions allows the diversity (given by Equation 1) to increase supra-exponentially, rather than linearly, with the number of elements *n*.

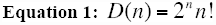

With only 10 fragments, the diversity reaches > 3×10^9^ unique sequences. Importantly, the equilibrium state of this architecture, as the number of recombination events *m* approaches infinity, is an equal distribution of all potential unique sequences (16). In practice, *m* will reach some finite number that may result in cassette biases. However, simply extending the cassette by one fragment can compensate for any modest biases of this architecture. Moreover, because the barcodes are made of a small number of known sequence fragments, reconstruction of barcode sequences even from highly error prone sources becomes possible. The order and orientation of each fragment within the cassette after recombination can be determined simply by performing *2n* pairwise alignments (each fragment in both orientations is aligned to the recovered sequence). This results in barcodes that are robust to many classes of sequencing errors.

**Figure 2.**
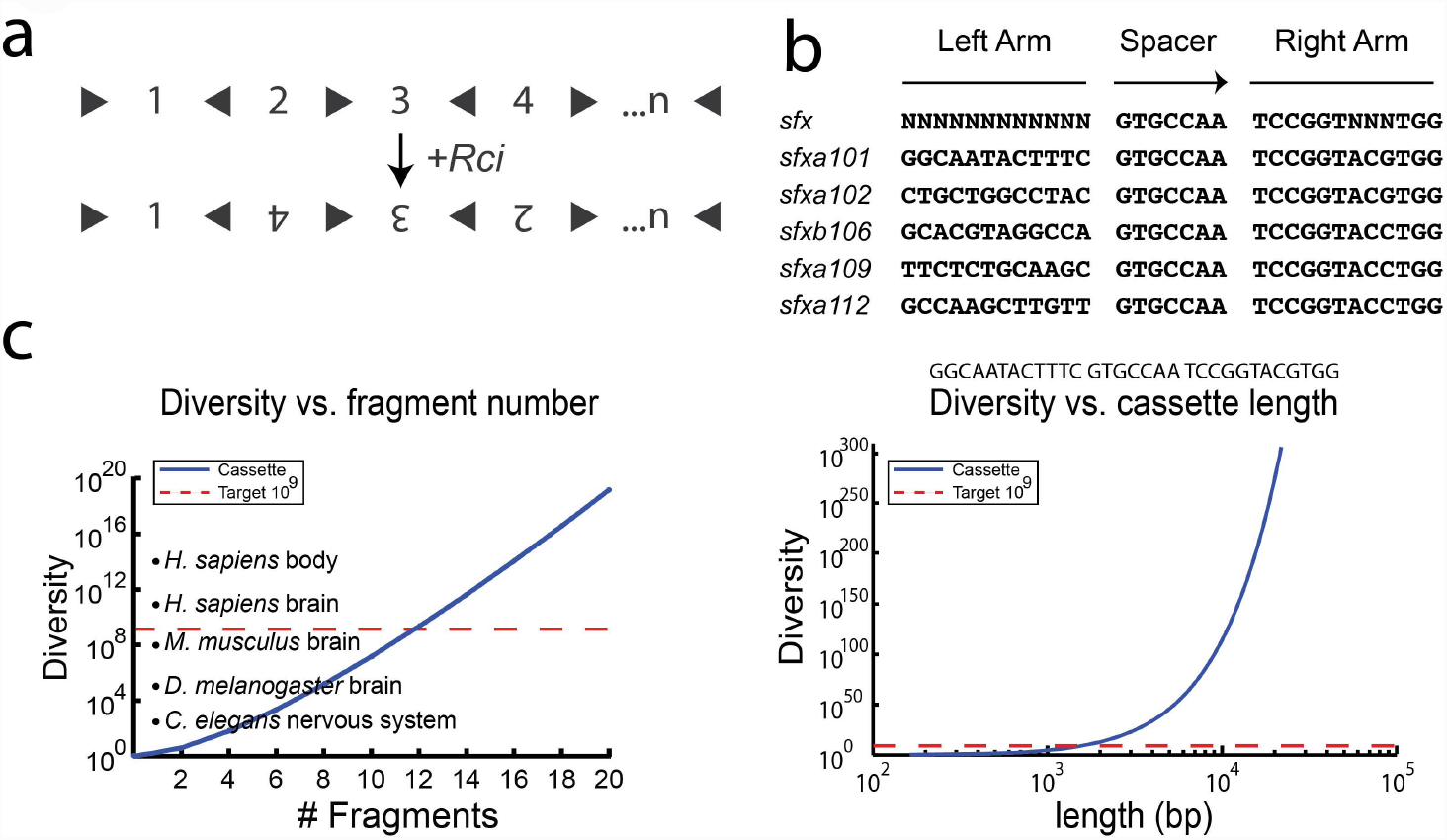
Design of Rci cassettes. **(A)** The Rci cassette is formed by concatenation of *n* sequence fragments, in which individual fragments are separated by a single *sfx* site - the orientation of which alternates between successive fragments. Upon Rci recombination, the cassette is shuffled. Because Rci cannot mediate excision events, the cassette remains the same length, but the relative position of each fragment within the cassette can change. **(B)** *sfx* sites share a central spacer sequence and right arm sequence, but have little sequence conservation in the left arm. 5 different *sfx* sequences were used in this study. **(C)** The diversity of the Rci cassette as a function of the number of fragments, *n.* Cassettes with >20 fragments can generate sufficient diversity to label all cells in complex organs/organisms. **(D)** The diversity as a function of length for Rci cassettes (assuming 100bp fragments). Lengths ∼1-2kb are needed to reach the requisite diversity of 10^9^.

We thus adopted a strategy based on DNA invertases - recombinases that can mediate only inversions (17). Rci (recombinase for clustered inversion) is a site-specific recombinase (SSR) of the integrase (Int) family, of which Cre is also a member (8). Rci recognizes 31bp *sfx* sites and mediates recombination events only between sites in inverted orientation (inversions) - it cannot mediate excision events between sites in the same orientation. Unlike other inversion systems, such as Hin and Gin (18), Rci does not appear to require any co-factors or enhancer sequences (19). Because of this, we selected Rci as our recombinase and designed a new barcode cassette in which segments of DNA are shuffled by inversion-only recombination events (Figure 2a).

### Design and synthesis of a 5BC cassette

Initially, we planned to synthesize DNA in which *n*=5 fragments, each flanked by *sfx* sites in opposite orientation, were concatenated to form a barcode cassette. After concatenation, each fragment is separated from its immediate neighbor by two *sfx* sites in opposite orientation (similar to the architecture proposed for Cre in Figure 1a). However, DNA synthesis constraints and plasmid stability forced us to redesign the cassette to minimize the effects of repetitive elements (the *sfx* sites). Ultimately, we employed an architecture in which individual fragments are separated by a single *sfx* site - the orientation of which alternates between successive fragments (Figure 2a), relying on several compatible *sfx* site variants to further reduce complexity (20) (Figure 2b).

This architecture achieves a somewhat lower diversity. Fragments originating in odd positions within the cassette (i.e. position 1, 3, 5, etc.) can only occupy odd positions after recombination. Likewise, fragments originating in even positions can only occupy even positions after recombination. This leads to a reduced diversity, given by Equation 2.

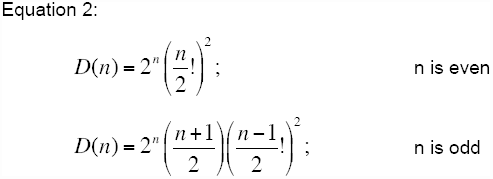

where *D* is the total diversity and *n* is the number of fragments in the cassette. Despite the reduction of diversity due to the modified architecture, only 12 fragments are required (*n*=12) to achieve our target diversity of >10^9^. Additional segments greatly increase the diversity making this a scalable approach (Figure 2c). Moreover, unlike the Cre-based architecture, the Rci-based cassettes reach our requisite diversity at reasonable cassette lengths of ∼1-2kb (Figure 2d).

**Figure 3.**
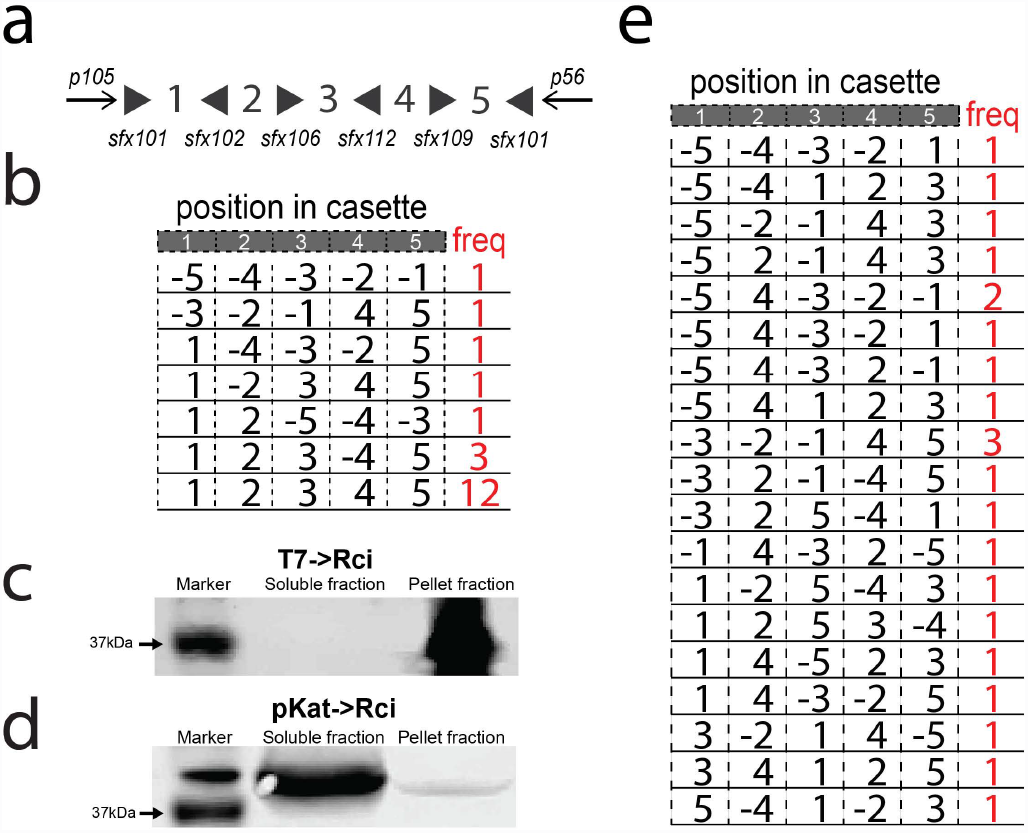
*In vivo* testing of Rci recombination on barcode cassettes. **(A)** A barcode cassette containing 5 fragments, which were separated by various *sfx* sites in alternating orientations, was synthesized. Anchor sequences flanked the cassette for use in PCR and sequence reconstruction. **(B)** Induction of Rci expression from the T7 inducible promoter results in shuffling in some, but not all cassettes. **(C)** Induction of Rci expression from the T7 inducible promoter results in Rci protein that is largely insoluble. **(D)** Expression of Rci from a medium-strength, constitutively active promoter results in expression of soluble Rci protein and **(E)** robust shuffling of the cassette.

Ultimately, a 5-fragment (100bp fragments) barcode cassette was synthesized utilizing 5 different *sfx* site sequences to decrease the repetitive nature of the cassette to aid in synthesis and replication. In addition, known anchor sequences (ANCHOR105 and ANCHOR56) positioned at either end of the cassette, were added outside of the recombination region to aid in sequence reconstruction. The final 5-fragment cassette, 5BC (Figure 3a), was cloned into a low-copy plasmid containing the Rci gene (resulting in plasmid IDP205). This ensures that all barcode cassettes that are transformed into bacterial cells will be exposed to the Rci coding sequence. Plasmid IDP205 (T7->Rci; 5BC) contains the Rci gene driven by the inducible T7 promoter. This plasmid is remarkably stable, showing no signs of recombination in the absence of induction across many generations (Supplementary Figure 2).

### Testing of the 5BC cassette *in vivo*

We transformed two populations of BL21-DE3 (NEB) bacterial cells with plasmid IDP205 (T7->Rci; 5BC) and grew 10mL cultures overnight. One culture was grown under conditions that induce the expression of Rci from the T7 promoter (see methods). After growth, cells were plated for clonal analysis on plates that did not support Rci expression. Twenty colonies were chosen for each condition (+ and - Rci induction) and analyzed by Sanger sequencing. Without induction of Rci expression, no recombination was observed (0 of 20 colonies sequenced, data not shown). Moreover, the induction of Rci led only to modest recombination - shuffling the cassette in 8 of the 20 reconstructed barcode sequences (Figure 3b). Interestingly, each of the final products could be explained by a single recombination event (Supplementary Figure 3).

We reasoned that the inefficient shuffling might be due to protein aggregation and insolubility due to high overexpression, as is often the case with T7 overexpression (21). To test this we fused an HA-tag at the N-terminus of Rci and tested the expression level and solubility via western blot. Indeed, we found that HA-Rci was found only in the insoluble fraction (Figure 3c), perhaps explaining the inefficient shuffling observed. Thus, we tested expression of HA-Rci from a different promoter, a medium strength constitutively active promoter, pKat (Registry of Standard Biological Parts: BBa_I14034), and found that the protein was soluble when expressed from this promoter (Figure 3d).

Therefore, we cloned the pKat promoter in place of T7 to make plasmid DIG35 (pKat->Rci; 5BC) and tested for shuffling efficiency. Briefly, we transformed DIG35 (pKat->Rci; 5BC) into Top10 competent cells and grew cultures overnight. To stop shuffling, plasmid DNA was isolated and the sequence for the Rci gene was removed via restriction enzyme digestion. Plasmids were retransformed and colonies were selected for Sanger sequencing. DNA was isolated from each colony and subjected to Sanger sequencing. Sequence reads were analyzed with our alignment algorithm in order to reconstruct full barcodes. Reads that could not be fully reconstructed from sequencing data were discarded from further analysis.

Expression of Rci from this promoter, pKat, resulted in robust shuffling (Figure 3e). Of the 22 reconstructed (3 sequences failed reconstruction) cassette sequences, each had undergone shuffling to yield 19 unique barcode sequences. Moreover, many of the final barcode sequences could only be explained by multiple recombination events. Based on these positive preliminary results, we next subjected the 5BC cassette to high throughput DNA sequencing.

### High throughput sequencing of shuffled 5BC

Advances in high throughput sequencing (HTS) have allowed for unprecedented access to massive quantities of DNA sequence data. We took advantage of HTS to sequence the shuffled 5BC cassettes at depths that allowed for a more thorough analysis of the actual *in vivo* behavior of barcode generation by Rci. Because of the length of our potential cassettes (∼1-2kb to reach diversities of 10^9^), we chose the PacBio sequencing platform.

Briefly, we transformed Top10 (Invitrogen) bacterial cells with either plasmid IDP205 (T7->Rci; 5BC - negative control) or DIG35 (pKat->Rci; 5BC) and allowed cultures to grow overnight. DNA was isolated and digested to release the barcode cassette. The barcode cassettes were then prepared for high throughput sequencing on the PacBio RS II and sequenced.

**Figure 4.**
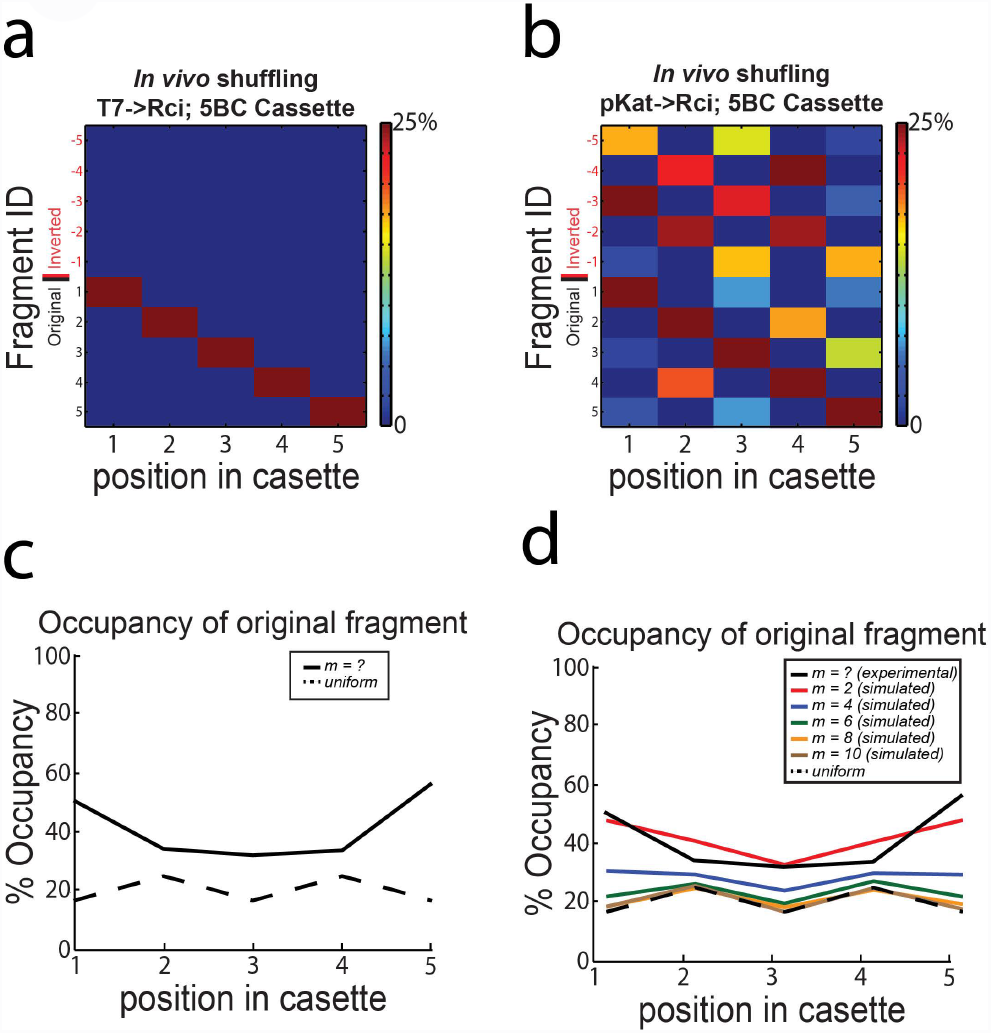
High throughput sequencing of 5BC Rci cassette after *in vivo* shuffling. **(A)** Cassettes are stable in the absence of Rci expression and can be reconstructed from PacBio sequencing data. **(B)** Expression of Rci from the constitutively active pKat promoter results in robust shuffling of the 5BC cassette. The colourmaps show the distribution of fragment occupancy at each position in the cassette. Colours are scaled from 0 to 25% for visualization. **(C)** The bias at each position was calculated as the number of times the original fragment appeared in its original position divided by the number of cassettes. The dotted black line indicates the expected occupancy of the original fragment at each position in a cassette with completely random occupancy. **(D)** The bias at each position within a cassette subjected to *m=2, 4, 6, 8, or 10* recombination events was simulated. The dotted black line indicates the expected occupancy of the original fragment at each position in a cassette with completely random occupancy. The solid black line indicates the data observed *in vivo.*

Using our algorithm, we reconstructed 5887 barcodes from 7203 circular consensus reads (see Methods) obtained from cells in which Rci was not expressed (plasmid IDP205; T7->Rci, 5BC). Of the reconstructed barcodes, 5886/5887 gave the original sequence (Figure 4a). These data indicated that the PacBio sequencing platform could handle the highly repetitive nature of the barcode cassettes and would allow for high throughput sequencing of cassettes without the introduction of recombination during sequencing from template switching or other sources. When Rci was expressed off of the pKat promoter (DIG35) the cassette was shuffled robustly (Figure 4b). Here, we reconstructed 5243 barcodes from 6105 circular consensus reads, of which there were 203 unique sequences. After shuffling, each position along the cassette is populated with a relatively even distribution of all of the possible fragments, with only slight biases at the ends of the cassette (Figure 4b). In other words, the occupancy at each position of the cassette, while still preferential for the original fragment sequence, approaches randomness (Figure 4c). This important observation indicates that there are no biological constraints on our design that prohibit the full exploration of the barcode space. Simulations of shuffling of a 5BC cassette under simple assumptions (see Methods for details) show that the occupancy at any given position reaches equilibrium after ∼6 or more recombination events per cassette (Figure 4d). Comparison of the simulated data and the data collected from *in vivo* shuffling suggests that our cassettes likely experienced 2-4 recombination events on average *in vivo* (Figure 4d).

Even this limited amount of recombination events resulted in an observed 203 unique sequences (out of a theoretical 384. *n=5, diversity =* 2^5^×3×(2!)^2^). Intuitively, after a small number of recombination events, the cassette remains biased at each position to its initial state (Figure 4d). As the number of recombination events increases, however, the cassette goes to equilibrium, and there is a nearly uniform distribution of each fragment at each position in the cassette (Figure 4d). In the limit, the cassette will reach an equilibrium state in which every possible barcode is equally probable (16).

This proof of principle experiment shows that *in vivo* recombination by Rci on a cassette is efficient at shuffling the original sequence into a unique barcode. However, the theoretical diversities of the 5BC cassette are well below our initial goals. Therefore, we sought to expand the cassette to achieve higher diversities.

### High throughput sequencing of shuffled 11BC

**Figure 5.**
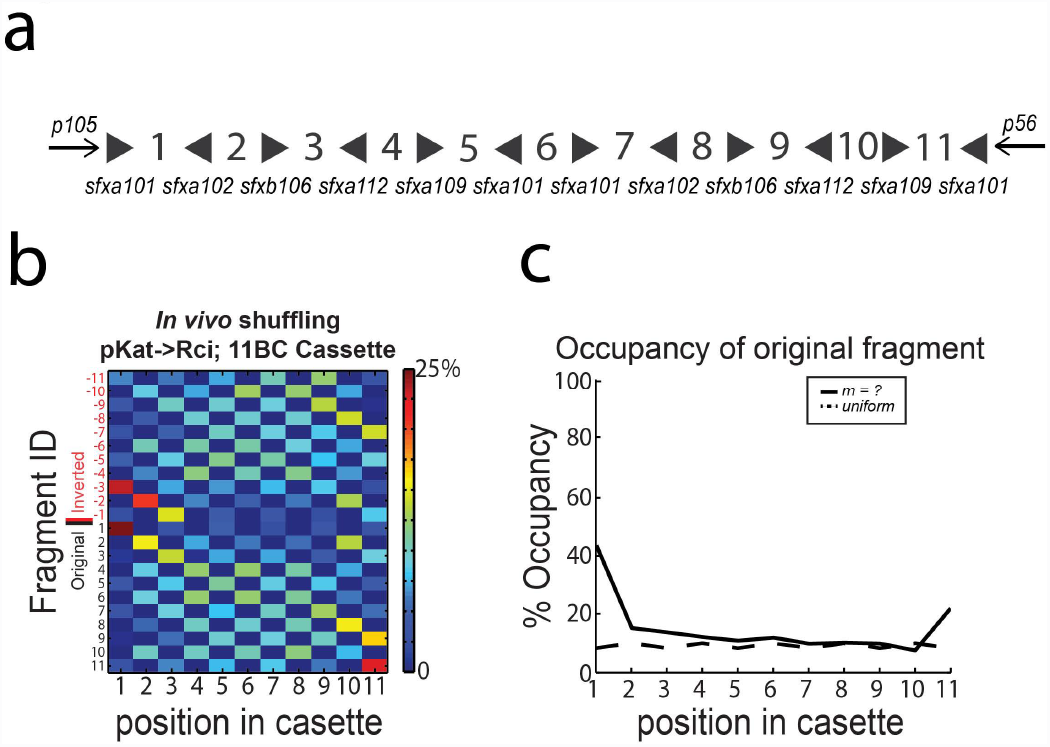
High throughput sequencing of 11BC Rci cassette after *in vivo* shuffling. **(A)** A 6-fragment extension was concatenated with the 5BC cassette to yield the 11BC cassette. **(B)** Expression of Rci from the constitutively active pKat promoter results in robust shuffling of a high-diversity 11BC cassette. The colourmaps show the distribution of fragment occupancy at each position in the cassette. Colours are scaled from 0 to 25%. **(C)** The bias at each position was calculated as the number of times the original fragment appeared in its original position divided by the number of cassettes. The dotted black line indicates the expected occupancy of the original fragment at each position in a cassette with completely random occupancy.

To explore the feasibility of achieving diversities that are capable of labelling large populations of cells, we expanded the cassette to 11 fragments. We synthesized a 6-fragment extension to our original cassette and concatenated this with our original 5BC cassette to create an 11BC cassette (Figure 5a). Importantly, plasmids harbouring this cassette were stable across many generations and showed no evidence of recombination in the absence of Rci expression (Supplementary Figure 4).

The theoretical diversity of this cassette is 2^11^×6×(5!)^2^ = 176,947,200. Unfortunately, there is currently no sequencing technology that has both the requisite depth and read-length to appropriately cover the potential diversity of the 11BC cassette. Nevertheless, we used high-throughput sequencing on the PacBio platform to sample the barcodes produced by the recombination of the 11BC cassette. Briefly, we transformed Top10 bacterial cells with plasmid DIG71 (pKat->Rci; 11BC cassette) and cultured the cells overnight. The barcode cassette was released by restriction digestion and subjected to high throughput sequencing on the PacBio RS II.

Shuffling of the 11BC cassette *in vivo* was efficient (Figure 5b). We reconstructed 1786 barcodes, of which 1723 were unique. Again we observe that, after shuffling *in vivo*, each position along the cassette is populated with a relatively even distribution of all of the possible fragments (Figure 5b), approaching a completely random cassette (Figure 5c). As suggested by simulations, further shuffling will lead to an increasingly random cassette (Supplementary Fig 5).

## DISCUSSION

To our knowledge, this is the first example of an *in vivo* barcoding scheme with the potential to scale to uniquely label all of the individual cells of an entire tissue or organism. The system, which takes advantage of recombinases to shuffle fragments within a cassette has several advantageous characteristics. Recombinases similar to Rci (i.e. Cre, FLP, PhiC31) have been successfully employed across a wide variety of organisms, suggesting that our barcoding system could be easily ported to organisms beyond bacteria, either by expressing Rci or alternatively, by employing designs that take advantage of asymmetric mutant recombination sites (22) of other recombinases. In addition, the system is highly scalable - addition of a single fragment to a cassette results in an exponential gain in diversity. Moreover, because the input space is small (only a handful of unique segments), each segment can be designed to be maximally orthogonal, thus rendering barcode readout highly robust to DNA sequencing errors. We took advantage of this fact in designing our barcode reconstruction algorithm, which relies on local alignment between the known input fragments and the final imperfect sequence recovered from high throughput sequencing.

In its current form, however, this paradigm has at least several shortcomings that will need to be addressed before the system can be used at a larger scale. First, the expression of the Rci protein must be controlled in order to induce shuffling at a specific time point and then stopped to prevent further recombination events. In theory this can be accomplished through the use of an inducible promoter (i.e. T7 promoter, Tetracycline-responsive promoter, etc.). In practice, the expression level and recombination efficiency will need to be monitored at different levels of induction to permit solubility, genomic stability, and optimal recombination kinetics. Second, the cassettes, despite their design, were still subject to various poorly understood biases. Further work will be needed on Rci (and other recombinases) to determine the pair-wise efficiencies between different recombination sites, the efficiency of recombination as a function of length between recombination sites, and to increase the recombination kinetics to achieve the maximal number of recombination events per unit time. As we saw in our simulations, a higher number of recombination events leads to far greater diversity and less bias. The biases that we detected, particularly in the case of the 5BC cassette, were likely exacerbated by the fact that we introduced a homogenous barcode into exponentially dividing cells, where the kinetics of cell division likely outpace that of recombination (at least initially). In terminally differentiated cells, such as neurons, this is less of an issue as the expression of the recombinase can be sustained for long periods of times (i.e. weeks or months). However, in the case of dividing cells, recombinase expression must be pulsed for short durations to allow shuffling and then abruptly stopped to permit lineage tracing. An additional limitation of our design is that the barcodes are carried on extrachromosomal plasmids and thus cannot be used for lineage tracing. Introduction of the cassette into the genome of bacteria can be accomplished by homologous recombination and should be straightforward. Finally, as described, this method relies on long sequence reads - which were obtained here by Sanger sequencing and/or high-throughput sequencing with the Pacific Biosciences platform. To achieve the orders of magnitude more reads necessary to track cells in complex populations will require the use of next-generation sequencing platforms with greater read-depth. Currently, Pacific Biosciences’s sequencing platform allows for ∼100k reads of >1kb. Alternative platforms, which produce a far greater number of reads per run (i.e. Illumina), lag behind in terms of read length but are consistently improving. A 3× increase in read length on the Illumina platform (from PE250 to PE750) would be enough to sequence our current cassettes and is likely to be possible in the near future. Alternatively, because the fragment sequences are known, it may be possible to generate subreads for each Illumina cluster by priming with various fragment sequences. These could then be assembled into full-length barcode reads. In addition, further research on Rci could allow for the optimization of barcode fragment lengths to permit recombination with shorter fragments - thereby decreasing the length of the cassette to be within reach of current next-gen sequencing platforms.

We have shown that *in vivo* shuffling of a cassette of DNA fragments by a recombinase - Rci - can generate significant diversities for cellular barcoding purposes. Barcoding of individual cells within bacterial or yeast populations should prove to be a useful tool for population geneticists and evolutionary biologists. The system has few moving parts (all that is needed is Rci and a barcode cassette) and is likely to work across a variety of higher organisms with optimization. This system, applied in other organisms could pave the way to the dissection of complex developmental programs, study of heterogeneity within tissues, and/or probing of interactions between cells in a population. In vivo barcoding will also pave the way to new explorations in systems biology, from high-throughput monitoring of non-cell-autonomous spread of genetic material to the variability of single-cell transcription profiles. In combination with other molecular tools, in vivo barcoding has the potential to provide biologists with unprecedented knowledge of the complex orchestration of single cells within populations.

## ACKNOWLEDGEMENTS

The authors would like to acknowledge Barry Burbach, Hassana Oyibo, Huiqing Zhan, Justus Kebschull, and Daniel Starer Stor for technical support. Additionally, we thank Josh Dubnau, Alex Koulakov, Ton Schumacher, and Reid Johnson for useful discussions.

